# The brain’s hedonic valuation system’s resting-state connectivity predicts weight loss and correlates with leptin

**DOI:** 10.1101/2020.01.27.921098

**Authors:** Liane Schmidt, Evelyn Medarwar, Judith Aron-Wisnewsky, Laurent Genser, Christine Poitou, Karine Clément, Hilke Plassmann

**Author notes:** Authors contributed equally and are listed in reverse alphabetical order (Evelyn Medarwar, Judith Aron-Wisnewsky). Authors contributed equally and are listed in reverse alphabetical order (Christine Poitou, Karine Clément). Author Contributions: HP, KC, CP and JAW conceived the project, and HP designed this study. JAW, CP and KC coordinated clinical investigation (MICROBARIA and LEAKY GUT). JAW, CP and KC contributed to the recruitment of participants with obesity involved in the bariatric surgery program. LG performed the surgery. LS analyzed data; EM assisted with data analysis under LS’s and HP’s supervision. HP and LS wrote a first draft of the manuscript, and all authors contributed to the final text.

## Abstract

Weight gain is often associated with the pleasure of eating foods rich in calories and lack of willpower to reduce such food cravings, but empirical evidence is sparse. Here we investigated the role that connectivity within the brain’s hedonic valuation system (BVS, the ventral striatum and the ventromedial prefrontal cortex) at rest plays (1) to predict weight gain or loss over time and (2) for homeostatic hormone regulation. We found that intrinsic connectivity within the BVS at rest (RSC) predicted out-of-sample weight changes over time in lean and obese participants. Counterintuitively, such BVS RSC was higher in lean versus obese participants before the obese participants underwent a drastic weight loss intervention (Roux-en-Y gastric bypass surgery, RYGB). The RYGB surgery increased BVS RSC in the obese after surgery. The obese participants’ increase in BVS RSC correlated with decreases in fasting state systemic leptin, a homeostatic hormone signalling satiety that has been previously linked to dopamine functioning. Taken together, our results indicate a first link between brain connectivity in reward circuits in a more tonic state at rest, homeostatic hormone regulation involved in dopamine functioning and ability to lose weight.

**Significance statement:** With obesity rates on the rise, advancing our understanding of what factors drive people’s ability to lose and gain weight is crucial. This research is the first to link what we know about the brain’s hedonic valuation system (BVS) to weight loss and homeostatic hormone regulation. We found that connectivity at rest (RSC) within the BVS system predicted changes in weight, differentiated between lean and obese participants, and increased after a weight loss intervention (gastric bypass surgery). Interestingly, the extent to which BVS RSC improved after surgery correlated to decreases in circulating levels of the satiety hormone leptin. These findings are the first to reveal the neural and hormonal determinants of weight loss, combining hedonic and homeostatic drivers of (over-)eating.

## Introduction

In Western societies today, more than half of adults are overweight or obese, and obesity rates are projected to continue to grow (OECD 2017). Despite the prevalence and severity of obesity, its neurobiological underpinnings and how they are changed upon weight loss in humans are not well understood.

Most previous cognitive neuroscience research has investigated differences in task-based activity between obese and lean participants using functional magnetic resonance imaging (fMRI). These studies found that exposure to high-calorie foods altered activity in brain regions involved in the hedonic aspects of food intake, such as reward and motivation processing (Rothemund et al. 2007; Stice et al. 2008; Volkow et al. 2008; Stoeckel et al. 2009), taste processing (Dagher 2007; Scharmuller et al. 2012) and cognitive control (Brooks et al. 2013; Pursey et al. 2014). In healthy participants, these brain systems were found to encode how much participants wanted to eat different foods (Plassmann et al. 2007, 2011) and to control potential cravings (Hare et al. 2009, 2011; Hutcherson et al. 2012).

Another stream of research has investigated tonic differences in the brain activity of obese and lean participants by capturing intrinsic connectivity among large-scale brain networks at rest. Studies in this area have found resting-state connectivity (RSC) differences in the salience, reward, default mode, prefrontal and temporal lobe networks (Coveleskie et al. 2015; Doornweerd et al. 2017; Garcia-Garcia et al. 2015; Kullmann et al. 2012; Wijngaarden et al. 2015) of obese and lean participants. Notably, RSC in these brain systems was shown to be altered by bariatric surgery-based weight-loss interventions (Li et al. 2018; Wiemerslage et al. 2017; Frank et al. 2014) shortly after those interventions (i.e., 4-12 weeks).

The goal of this paper is to put these different streams of research together and shed light on a potential link between RSC in the brain and changes in weight. We applied a theory-driven approach to investigate (1) differences in RSC in the brain’s hedonic valuation (i.e., the ventromedial prefrontal cortex [vmPFC] and striatum (Bartra et al. 2013)) and control systems (i.e., the dorso- and ventrolateral prefrontal cortex)(Hare et al. 2009, 2011; Hutcherson et al. 2012) between participants with severe obesity and lean control participants and (2) whether a longer term (i.e., 24 weeks) weight change due to bariatric surgery would affect RSC in these systems. We then used the changes in RSC in these regions to *formally* predict our lean and obese participants’ weight changes over time.

Bariatric surgery—specifically Roux-en-Y gastric bypass (RYGB)—serves as a unique and effective theoretical model for the questions of our work because it leads not only to rapid and major weight loss (needed for our formal prediction analysis to have the required variance) but also to improvements in hormone profiles involved in the homeostatic control of food intake (Sjöström et al. 2012; Abdennour et al. 2014). It thus allows us to move beyond previous correlational evidence and make quasi-causal links between RSC in the brain’s hedonic valuation system and hormonal homeostatic regulators of food intake.

A separate stream of clinical research has made major progress in understanding the neuronal circuitry involved in the control of energy homeostasis, including the role of bariatric surgery. For example, before surgery most obese individuals have extremely high fasting-state leptin levels, but the action of leptin to signal satiety is impaired (Myers et al. 2010). After RYGB surgery, levels of circulating leptin drop rapidly, and its ability to signal satiety improves (Faraj et al. 2003). Moreover, leptin has direct and indirect links to the brain’s hedonic valuation system. For example, it inhibits ventral tegmental area (VTA) dopamine neurons (Palmiter et al. 2007) that are known to directly project to the brain’s hedonic valuation system (Haber et al. 2003). Against this background the final goal of this paper was to explore why the surgery might alter RSC in the brain’s hedonic valuation system by exploring its links to surgery-induced changes in fasting-state serum leptin.

The contribution of this work is to provide first evidence that resting-state connectivity in the brain’s hedonic valuation system (1) predicts weight loss, (2) differs between lean and obese individuals, (3) is altered by RYGB surgery and (4) is linked to RYGB surgery-induced changes in leptin levels (i.e., a marker of the hormonal homeostatic system of food intake control). This work is one of the first to integrate hedonic and homeostatic factors in food intake control (Berthoud 2006).

## Materials and Methods

### Experimental setup

The experimental procedure was conducted in accordance with the Declaration of Helsinki and received approval from the local ethics committee for obese participants and from INSERM for lean participants (Comités de Protection des Personnes [CPP], Ile-de-France). Informed written consent was obtained from all participants prior to study inclusion. The obese participants took part in the Microbaria and Leaky-gut protocols that are registered as clinical trials NCT01454232 (Microbaria) and <u>NCT02292121</u> (Leaky gut). The resting-state data presented in this paper was acquired as part of a multi-study project including different experimental tasks such as task-based fMRI, metabolic and faeces samples for microbiota analysis. The results of those other tasks are presented elsewhere (Aron-Wisnewsky et al. 2018); the focus of this paper is the differences in resting-state connectivity between lean and obese individuals and before and after bariatric surgery-induced weight loss. The scanning session consisted of a brief introduction and training, two task-based fMRI sessions, a structural MRI scan, and the resting-state fMRI scan presented in this paper.

Data were collected at two time points (T0 and T6) separated by six months. The participants with severe obesity underwent RYGB surgery shortly after their scanning session at T0; they were followed in the nutrition department at the Specialized Obesity Centre for Obesity and Obesity Surgery at Pitié-Salpêtrière Hospital in Paris. MRI data was collected at the Centre for Neuroimaging (Cenir) at the Institut du Cerveau et de la Moelle épinière (ICM) at Pitié-Salpêtrière Hospital in Paris. The lean participants’ brains were also scanned at the same facilities of the Cenir twice to control for the effect of time.

### Participants

A total of 64 female participants were enrolled at T0, including 45 lean participants and 19 with severe obesity. We recruited only female participants in an attempt to keep gender influences constant^43^. Additional standard fMRI inclusion criteria were right-handedness, normal to corrected-to-normal vision, no history of substance abuse or any neurological or psychiatric disorder, and no medication or metallic devices that could interfere with performance of fMRI. The participants with obesity and the lean controls were recruited based on their body mass index (BMI), which was on average 22 ± 0.3 kg/m^2^ for the lean participants and 45 ± 1 kg/m^2^ for the candidates for bariatric surgery with severe obesity in agreement with international guidelines (see Tables 1 and 2 for more details on clinical characteristics and body composition).

**Table 1:**
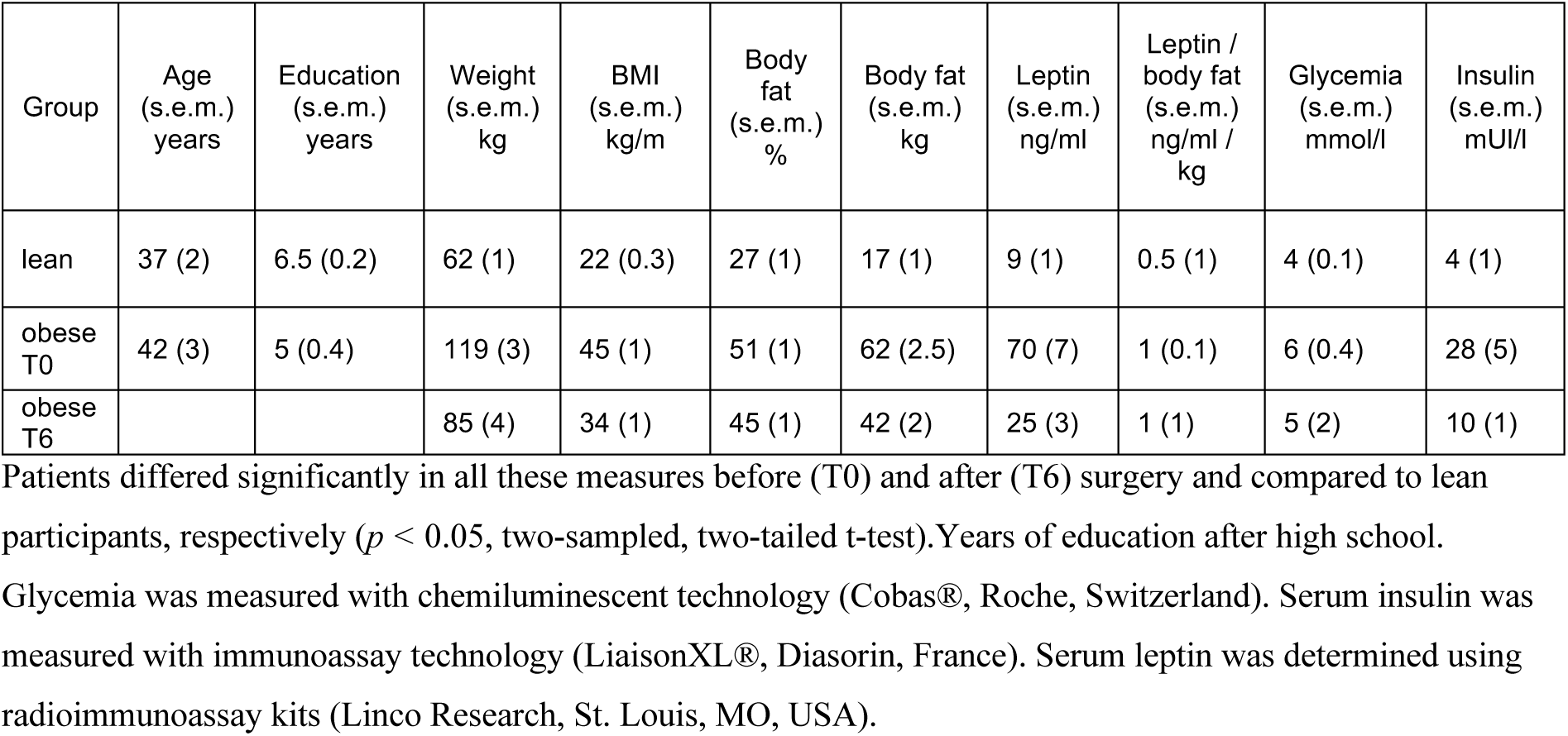
Participant main characteristics.

**Table 2:** Main effect of group on vmPFC resting state connectivity.

| Obese > Lean participants at T0 |  |  |  |  |  |  |
| --- | --- | --- | --- | --- | --- | --- |
| Region | BA | size | x | y | z | Peak z-score |
| Cerebellum |  | 729 | -20 | -80 | -44 | 5.30 |
|  |  | 848 | 14 | -78 | -44 | 4.95 |
| dIPFC | 47 | 166 | -34 | 10 | 46 | 4.26 |
| Obese < Lean participants at T0 |  |  |  |  |  |  |
| Region | BA | size | x | y | z | Peak z-score |
| Ventral striatum |  | 172 | -10 | 10 | -4 | 4.51 |
| Hippocampus |  | 283 | 22 | -48 | 8 | 4.44 |
| Obese > Lean participants at T6 |  |  |  |  |  |  |
| Region | BA | size | x | y | z | Peak z-score |
| IFG | 45/46/47 | 243 | -54 | 38 | 12 | 4.74 |
| vIPFC | 47/11 | 191 | 44 | 42 | -12 | 4.15 |
|  | 10/11 | 228 | 30 | 60 | 0 | 4.08 |

This table reports the peak coordinates and z-score values for lean participants compared to obese patients before at T0 and six months after bariatric surgery at T6. All peaks surpassed a voxel-wise threshold of $p_{FDR} < 0.05$ false discovery rate (FDR) corrected on the cluster level. The xyz coordinates correspond to the Montreal Neurological Institute (MNI) space. dIPFC: dorsolateral prefrontal cortex; IFG: inferior frontal gyrus; vIPFC: ventrolateral prefrontal cortex.

Of the 64 individuals recruited for the study, 20 participants were excluded before starting our analyses due to the following predefined exclusion criteria: two lean and three participants with obesity were excluded because of extensive head motion (≥3.5 mm), 10 lean participants were excluded because they did not return for their six-month MRI evaluation, and three lean and two obese participants had incomplete rfMRI data. Therefore, a total of 44 (30 lean and 14 obese) participants were included in all analyses concerning within-participant time effects (e.g., T0 versus T6) and group by time interactions. Note, we could still perform analyses concerning between-participant group effects (e.g., obese versus lean) at baseline (T0) for 56 participants (40 lean, 16 obese) who had available data at T0.

### Roux-en-Y gastric bypass (RYGB) surgery

Roux-en-Y gastric bypass surgery is a surgical intervention reserved for the most severe forms of obesity (BMI ≥ 40kg/m^2^ or BMI ≥ 35kg/m^2^ with obesity-related comorbidities) (44). RYGB creates a small gastric pouch directly linked to the distal small intestine with a gastro-jejunal anastomosis. The remaining part of the stomach and the proximal small intestine are bypassed, creating a Y-Roux limb (see 45 for details). The resulting Y-shaped gastric bypass, called the Roux limb, replaces most parts of the stomach and the first section of the small intestine, the duodenum. Ingested food thus directly goes from the newly created gastric pouch to the small intestine, which reduces the nutrients and calories absorbed from food.

In our study, the RYGB surgery was performed laparoscopically. All participants were clinically assessed before and one, three and six months post-surgery, as recommended by international guidelines(Fried et al. 2014). The clinical assessments included obesity-related diseases and anthropometric measures estimated by whole-body-fan-beam dual-energy X-ray absorptiometry (DXA) (Hologic Discovery W, software v12.6, 2; Hologic, Bedford, MA, USA), as detailed in Ciangura et al. 2010. Variables included weight, body mass index (BMI) and total body fat in kg and percent (Table 1).

### Blood hormone sampling

Blood samples were collected once from the lean participants (at T0), and twice for participants with obesity before (T0) and six months after RYGB (T6). Venous blood samples were collected in the fasting state (12-hour fasting) for determination of glycemia, insulinemia and leptin. Glycemia was measured with chemiluminescent technology (Cobas®, Roche, Switzerland). Serum insulin was measured with immunoassay technology (LiaisonXL®, Diasorin, France). Serum leptin was determined using radioimmunoassay kits (Linco Research, St. Louis, MO, USA).

### Brain imaging data

#### Image acquisition

Resting-state fMRI scanning was conducted during a 10-minute scanning sequence after the participants took part in several task-based fMRI sessions. Participants were instructed to keep their eyes closed and to relax, but not to fall asleep.

T2*-weighted echo planar images (EPI) with BOLD contrast were acquired using a 3T Siemens Verio scanner. An eight-channel phased array coil was used to assess whole-brain resting-state activity with the following ascending interleaved sequence: Each volume comprised 40 axial slices, TR = 2s, TE = 24ms, 3-mm slice thickness; 0.3-mm inter-slice gap corresponding to 10% of the voxel size, FOV = 204 mm, flip angle = 78°. For each participant a total of 304 volumes were obtained. The first five volumes of the resting-state scan session were discarded to allow for T1 equilibrium effects. A single high-resolution T1-weighted structural image (MPRAGE) was acquired, co-registered with the mean EPI image, segmented and normalized to a standard T1 template. Normalized T1 structural scans were averaged across lean and obese participants respectively to allow group-level anatomical localization.

#### Preprocessing

Data was analysed using the Statistical Parametric Mapping software (SPM12; Wellcome Department of Imaging Neuroscience) along with the Functional Connectivity toolbox (CONN toolbox: www.nitrc.org/projects/conn, RRID: SCR_009550). Preprocessing in SPM included spatial realignment to estimate head motion parameters. This preprocessing step was done prior to slice-time correction, because slice-time correction can lead to systematic underestimates of motion when it is performed as a first preprocessing step (Drysdale et al. 2017). After realignment, preprocessing included the standard steps: slice-time correction, co-registration, normalization using the same transformation as structural images, spatial smoothing using a Gaussian kernel with full width at half maximum of 8 mm, and temporal band pass filtering between 0.01 and 0.1 Hz.

#### Nuisance signal removal

Nuisance signal removal was performed on the preprocessed time-series data with the CONNv16 toolbox; it included linear and quadratic de-trending to adjust for scanner drift, removal of nuisance signals related to head motion, and physiological variables by means of regression analyses. More specifically, the nuisance regression included 18 head motion parameters calculated during spatial realignment (roll, pitch, yaw and translation in three dimensions, plus their first and second derivatives), non-neuronal signals from eroded white matter (WM) and cerebral spinal fluid (CSF) masks, and regressors for outlier volumes. Individual WM and CSF masks were obtained by segmentation of each participant’s structural MPRAGE image into tissue probability maps using SPM12. The WM and CSF masks were further eroded to reduce partial volume effects. We used CONNv16’s ART-based function to identify outlier volumes with a global signal *z*-value threshold of 3 and an inframe displacement threshold of ≥ 0.5mm, corresponding to the most conservative setting in the CONNv16 toolbox (95th percentiles in normative sample). Note that the nuisance signal regression and band-pass filtering were performed simultaneously, only on volumes that survived head motion censoring. We used a rather lenient head motion threshold of ≥3.5 mm in order to not exclude our morbidly obese participants, who moved significantly more than the lean ones. After preprocessing, the smoothed residual time-series data, co-registered to MNI space, were used for the subsequent statistical analysis steps.

### Statistical analyses

We focused on a seed-to-voxel correlational analysis approach in order to investigate how functional connectivity between brain regions implicated in dietary decision-making and self-control is affected by obesity and bariatric surgery.

#### Seed region of interest (ROI)

Prior studies using fMRI have suggested that the vmPFC is a key region of the brain’s hedonic valuation system that encodes both expected and experienced value (Bartra et al. 2013). Previous work has shown that the vmPFC is activated under dietary decision-making (Plassmann et al. 2007, 2010) and self-control (Hare et al. 2009, 2011; Hutcherson et al. 2012), and individual differences in vmPFC anatomy are a marker for dietary regulatory success during dietary self-control (Schmidt et al. 2018). We therefore based the seed ROI on the vmPFC.

Specifically, the seed ROI was defined by the neurosynth (5.20.13) website using the “reverse inference” map for “vmPFC”. The mask was thresholded at *p* < .0001 uncorrected, after smoothing the Z-map with a 6mm FWHM kernel and averaging Z-scores across the left and right hemispheres to create a symmetrical map. We further resliced each mask to the lean controls’ and obese patients’ normalized mean EPI images to make sure that all voxels were within the vmPFC in our participant sample.

In order to investigate differences in the resting-state connectivity between lean and obese participants and between T0 and T6, a multiple regression analysis correlated the averaged BOLD signal from the vmPFC seed region of interest to the BOLD signal in each voxel of the brain for each participant. The Pearson’s r for each voxel was then transformed into a z-score using Fisher r-to-z transformations to obtain normally distributed functional connectivity (FC) coefficient maps. Individual FC coefficient maps were subjected to second-level random-effects factorial analysis of variance (2×2 ANOVA) crossing participant group (obese vs. lean participants) and time point (T0 vs. T6). We considered a false discover rate (FDR)-corrected significance threshold of *p*_FDR_ < 0.05 at the cluster level and further explored results at an uncorrected voxel-wise threshold of *p* < 0.001 to report the full extent of effects (Poldrack et al. 2008).

#### Out-of-sample cross-validation of the correlation between vSTR-vmPFC connectivity and weight loss

To test whether weight loss can be predicted from vSTR-vmPFC connectivity, we conducted the following leave-one-participant-out predictive analysis: First, z-values of vSTR-vmPFC functional connectivity were extracted for each participant and averaged across the voxels of the vSTR cluster that displayed an interaction effect. In other words, resting-state activity in these vSTR voxels correlated more strongly to vmPFC resting-state activity after surgery (at T6) compared to before surgery (at T0) in the obese compared to the lean participants. The average z-values for the vSTR cluster were then used to conduct 44 linear regressions that determined independent weights of the vSTR-vmPFC connectivity on weight loss (kg at T6 minus kg at T0) over 43 participants following equation i:

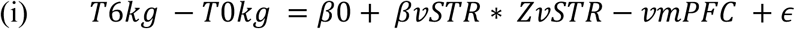

Each time, the weight (*βvSTR*) of the vSTR-vmPFC connectivity on weight loss obtained from 43 participants together with the z-value for vSTR-vmPFC connectivity extracted from the vSTR cluster in the left-out participant was regressed to predict weight loss for the left-out participant 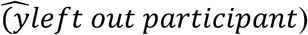 using the *glmval* function in matlab following equation ii:

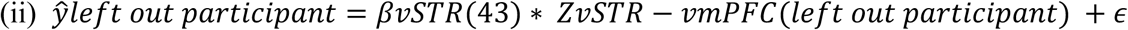

Last, we quantified the association between the predicted and observed levels of weight loss by using Pearson’s correlation that was tested for significance by using both parametric one-sampled t-tests and non-parametric permutation tests (1,000 permutations).

#### Hormone correlation analysis

We conducted correlation analyses to explore whether the changes due to the weight loss intervention in vmPFC-vSTR resting-state connectivity covaried with changes in leptin per kg body fat lost after surgery as a hormonal marker of homeostatic control of food intake. To this aim, Pearson’s correlation coefficient ρ was calculated following equation iii:

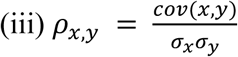

with *cov* corresponding to the covariance of *x* and *y* and σ corresponding to the standard deviation of *x* and *y*. Specifically, *x* corresponded to the change in raw serum leptin per kg body fat lost after surgery, according to equation iv:

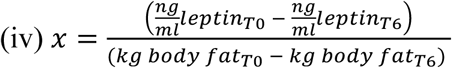

Because leptin is produced by white adipose tissue cells, the change in ng/ml leptin after surgery covaries significantly with kg body fat lost after surgery (Pearson’s ρ = 0.68, *p* = 0.007). We therefore considered the ratio as a measure of interest in order to account for the dependency between body fat and leptin. The ratio of the changes in leptin per kg body fat lost from T0 to T6 reflects the change of serum leptin levels per kg body fat lost after bariatric surgery. This ratio *x* was correlated to the change in vmPFC-vSTR connectivity after surgery *y. Y* was computed following equation v:

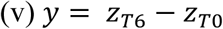

Mean connectivity values (*z*_*vmPFCtovSTR*_) were extracted for each obese participant at T0 and T6 from the ventral striatum cluster that displayed a significant connectivity to the vmPFC seed ROI for the interaction group (obese>lean) by time point (T6>T0) (MNI coordinates = [-10 6 -2], *p* < 0.001 uncorrected, extend threshold 50 voxels).

The significance of Pearson’s correlation coefficients was tested by conducting both parametric one-sampled t-tests and non-parametric permutation tests, which are less sensitive to individual outliers and estimated the 95% confidence intervals (CI) for correlations due to chance based on 10,000 permutations of the observed data.

#### Availability of materials and data

Code and data sets analysed in the current study are available from the corresponding authors on request.

## Results

We used resting-state magnetic resonance imaging to scan the brains of lean and obese participants (*n* = 64) twice, six months apart (see Table 1 for details). Importantly, the patients were scanned before and six months after undergoing RYGB surgery. We then analysed differences in the connectivity of the vmPFC – an important hub for dietary decision-making (Plassmann et al. 2007, 2010; Hare et al. 2009, 2011; Hutcherson et al. 2012) to other brain regions at rest. We sampled blood once in the lean participants (at the time of the first fMRI scan) and twice in the obese participants (pre- and post-RYGB surgery) to assess differences in serum leptin and how these differences were linked to changes in body fat and RSC connectivity in the obese patients before and after RYGB surgery.

### Differences in resting-state connectivity of the vmPFC in participants with obesity compared to lean participants

We first investigated differences in RSC in the brain’s hedonic valuation system with the vmPFC as seed between the obese and lean participants. In other words, we looked at the main effect of participant group irrespective of time^*^ and found that participants with obesity presented stronger vmPFC resting-state connectivity to a set of frontal brain regions including the dorsolateral prefrontal cortex (dlPFC), the ventrolateral prefrontal cortex (vlPFC) (cluster-corrected *p*_FDR_ < 0.05).

Post hoc comparisons between groups further revealed that at baseline (T0), severely obese patients compared to lean participants displayed stronger vmPFC connectivity to cognitive regulation nodes such as the dlPFC (Figure 1a, Table 2). After six months (T6), participants with obesity continued to have stronger vmPFC to vlPFC RSC (cluster-corrected *p*_FDR_ < 0.05; Figure 1b, Table 3).

**Fig. 1.**
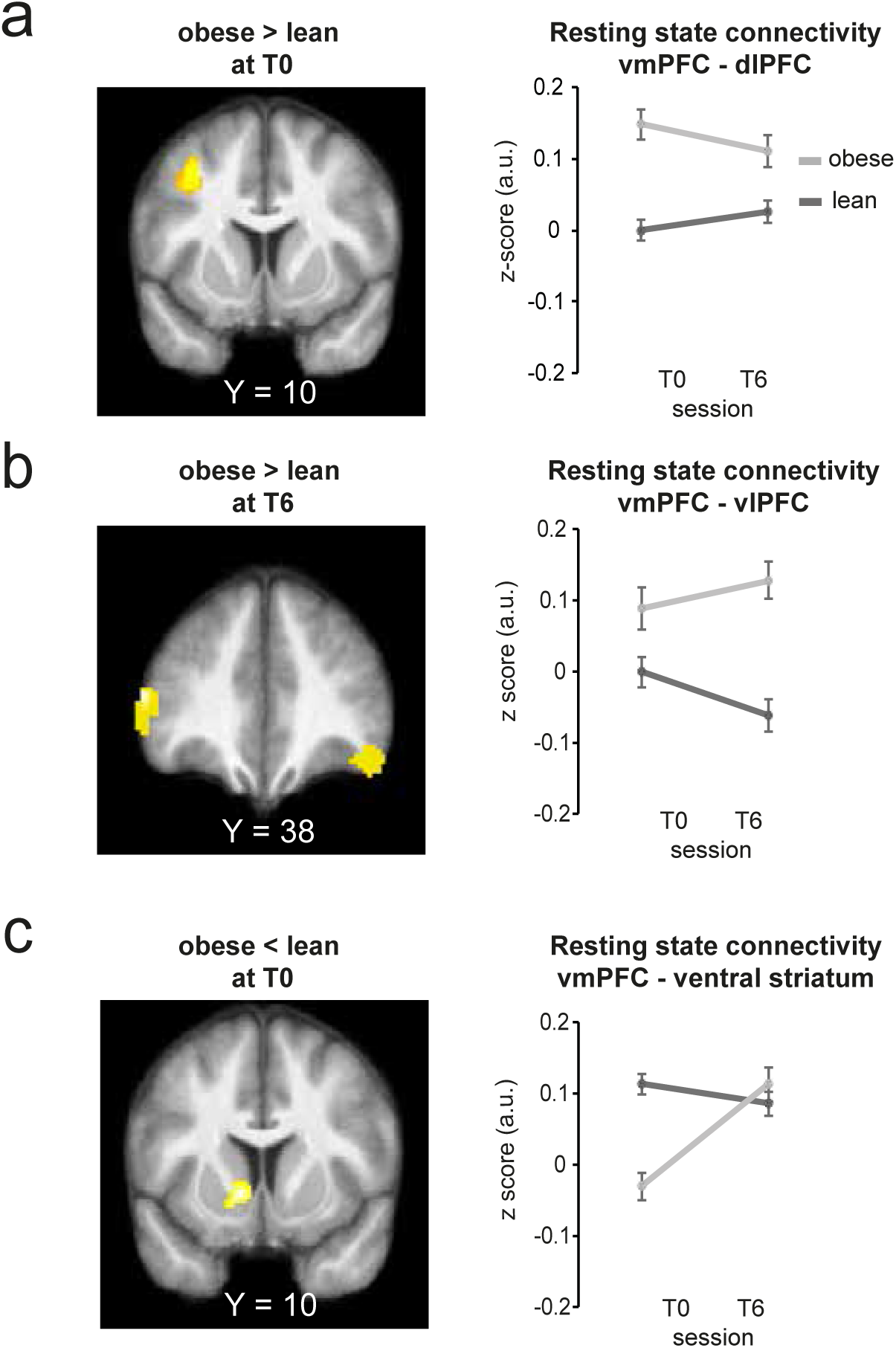
Comparisons of vmPFC to brain resting-state connectivity in lean and obese participants before and after bariatric surgery. SPMs of the seed-to-voxel resting-state connectivity between the vmPFC seed ROI and the rest of the brain, in obese > lean participants at **(A)** baseline (T0, *N* = 56, *p* < 0.001 uncorrected, extend threshold *k* = 166 voxels), **(B)** 6 months later (T6, *N* = 44, *p* < 0.001 uncorrected, extend threshold *k* = 191 voxels) and **(C)** in obese < lean participants (T0, *N*=56, *p* < 0.001 uncorrected, extend threshold *k* = 172 voxels). Significant voxels are displayed for visualization purposes in orange at *p* < 0.001 uncorrected, with an extend threshold *k* corresponding to a false discovery rate (FDR) corrected threshold of *p*_FDR_ < 0.05 on the cluster level for each contrast, respectively. SPMs are superimposed on the average structural image obtained from the lean participants. The [*x, y, z*] coordinates correspond to MNI coordinates and are taken at maxima of interest. The line graphs on the right of each SPM depict average correlation coefficients between resting state activity of the seed region, the vmPFC and the (a) dlPFC, (b) right vlPFC, and (c) ventral striatum at baseline (T0) and six months later (T6) in lean (dark grey) and obese (light grey) participants.

Another set of post hoc comparisons between groups showed weaker vmPFC connectivity to motivational nodes such as the ventral striatum (vSTR) (cluster-corrected *p*_FDR_ < 0.05; Figure 1c, Table 2) at baseline (T0). Interestingly, there were no differences between lean and obese participants in vmPFC-vSTR connectivity six months later (T6). Next we investigated the effect of surgery on RSC (i.e., the interaction between group and time).

### Effects of bariatric surgery on vmPFC connectivity

We investigated whether RYGB surgery affected the RSC of the vmPFC and, if so, whether it would affect its RSC to other brain regions involved in reward and motivation processing and control. In more detail, we compared the difference in the RSC of the vmPFC in the participants with obesity after versus before RYGB surgery to the change over time in the RSC of the vmPFC in the lean participants (i.e., the obese group > lean group by time T6 > T0 interaction). We found stronger RSC between the vmPFC and the vSTR RSC for this interaction (MNI coordinates [-10 6 -2], *p*_unc_ < 0.001, extend threshold *k* = 50 voxels; Figure 2a).

**Figure 2:**
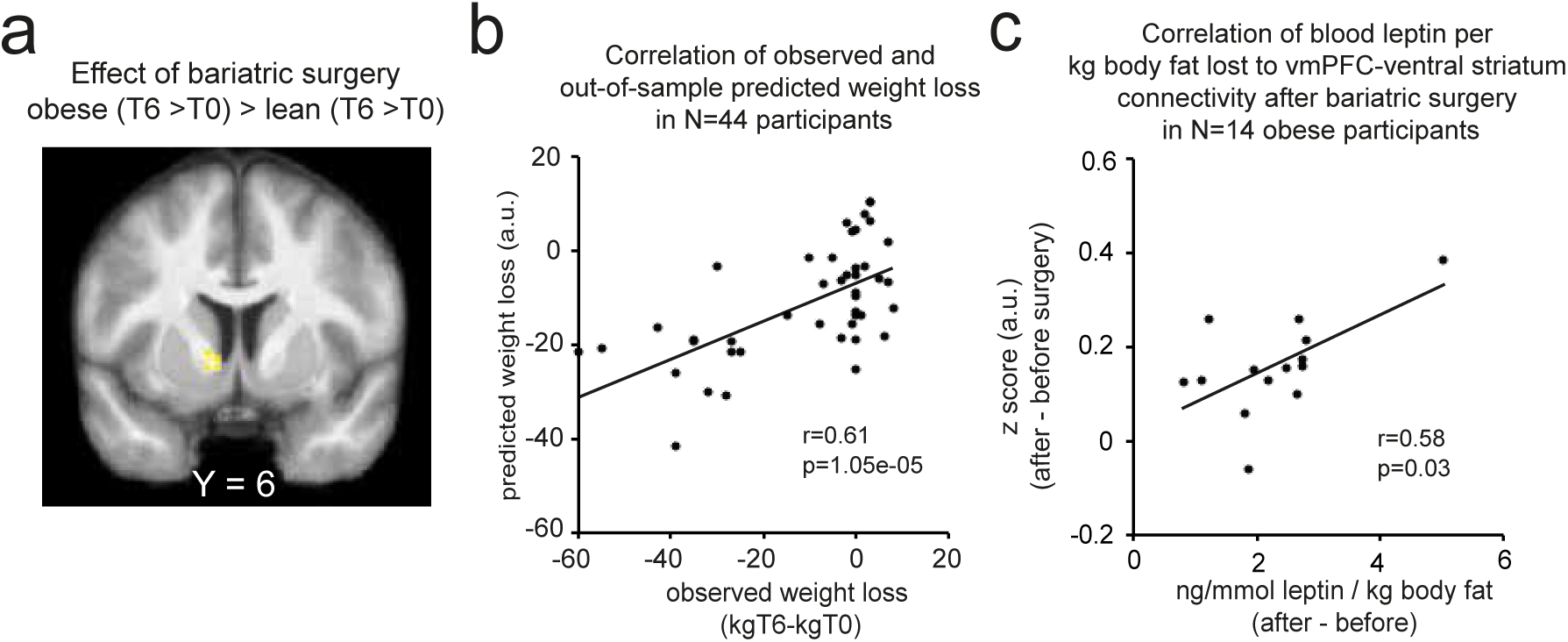
Effect of bariatric surgery on vmPFC to brain resting-state connectivity. **(A)** Resting-state activity in the vmPFC seed correlated significantly more to resting-state activity in the ventral striatum in obese participants after surgery compared to before surgery and to lean participants controlling for the time between baseline (T0) and six months later (T6) assessments (*N* = 44, *p* < 0.001 uncorrected, *k* = 50 voxels). SPMs are superimposed on the average structural image obtained from the lean participants. The [*x, y, z*] coordinates correspond to MNI coordinates and are taken at global maximum. **(B)** Scatterplots depict in all participants (*N* = 44) the correlation between observed weight loss (kg body weight at T6 minus kg body weight at T0) and predicted weight loss obtained from an out-of-sample cross-validation of the association between weight loss and vSTR-vmPFC connectivity. Dots correspond to obese participants. **(C)** Scatterplots depict in obese participants (*N* = 14) the change observed after, compared to before, bariatric surgery in vmPFC-ventral striatum resting state connectivity (average correlation coefficients) as a function of ng/mmol leptin per kg body fat lost. Dots correspond to obese participants.

### Out-of-sample prediction of weight loss over time across all participants

We then examined whether the changes in vSTR-vmPFC RSC could predict the changes in participants’ weight between the two time points using a leave-one-sample-out (LOSO) predictive analysis. When we based the prediction of weight loss on information about the vmPFC-vSTR RSC, there was a significant positive association between predicted and observed weight change (*r* = 0.61, *p* = 1.05e-05, 95% CI due to chance: -0.24–0.25; Figure 2b).

### Individual differences in relative fasting-state leptin determine changes in vmPFC-to-ventral striatum RSC

Finally, we explored how much the change in vmPFC-vSTR RSC after RYGB surgery was moderated by changes in serum leptin, taking into account the reduction of body fat. The adipose tissue secreted hormone leptin is well-known to contribute to signalling satiety and to stop food intake via inhibition dopamine receptors in the VTA and melanocortin (i.e., MC4) receptors in the hypothalamus. As expected, leptin and body fat were elevated in participants with obesity before surgery and decreased significantly post-surgery (% body fat: *t*(13) = 9.9, *p* < 0.001; kg body fat: *t*(13) = 13.7, *p* < 0.001 ng/ml leptin: *t*(13) = 5.6, *p* < 0.001; two-tailed, paired t-test; Table 1). When correlating the decrease in leptin per kg of body fat loss after RYGB surgery to the increase in vmPFC-vSTR resting-state connectivity after surgery, we found a significant positive correlation (Pearson’s ρ = 0.58, *p* = 0.03, 95% CI due to chance: - 0.46–0.46; Figure 2c). In other terms, participants with obesity who lost more circulating leptin per unit of fat mass post-RYGB were also those who had the most increased vmPFC-vSTR resting-state connectivity post-RYGB.

## Discussion

Our study provides first evidence using an out-of-sample prediction across all our participants that changes in RSC between the vmPFC and vSTR predicted how much weight participants lost over a period of six months. The vmPFC and the vSTR are two key regions within the brain’s hedonic valuation system involved in the processing of reward and motivation (Knutson et al. 2005; Rangel et al. 2008). Our finding is to the best of our knowledge the first to uncover an association between the propensity to lose weight over time and the connectivity of neural hubs at rest within the brain’s hedonic valuation system.

Interestingly, the RSC within the same system was attenuated in our obese participants when compared to the lean ones. This result parallels findings using task-based fMRI that showed a desensitization of the brain’s reward circuitry in response to food rewards in participants with obesity. Interestingly, such a desensitization has been described as similar to what happens in those who are addicted to drugs and other rewards (4). More specifically, several studies have shown that obesity shares some behavioural and neural similarities with drug addiction, such as overconsumption of certain types of highly palatable (HP) fat- and sugar-rich food, altered inhibitory control of food intake, and tolerance and withdrawal symptoms from HP food (Kable et al. 2007; Carter et al. 2016). On the neural level, these addictive behaviours have been linked to altered dopamine signalling in the brain’s reward system involving the vSTR and vmPFC (Carter et al. 2016; Volkow et al. 2012). Our results extend these links between obesity and diminished reward processing by showing that they might also be at play when participants are in a more general state of rest, affecting intrinsic connectivity in the brain.

Our study further found that a weight loss intervention based on bariatric surgery increased the vmPFC-vSTR. This finding suggests a reintegration of the brain’s hedonic valuation system in the obese participants after RYGB surgery to a level similar to that observed in the lean participants. Such a reintegration might be related to improved functioning of dopaminergic projections from the midbrain to regions of the brain’s hedonic valuation system. Our findings parallel those from positron emission tomography studies of dopaminergic functioning in patients with obesity. Specifically, dopamine D2 receptor availability has been shown to increase six weeks post-RYGB surgery (Steele et al. 2010), reaching levels similar to those observed in non-obese controls.

To strengthen the idea of a possible link between our findings and dopamine functioning, we found that vmPFC-vSTR RSC was positively correlated with the reduction of fasting-state serum leptin (taking into account fat-mass loss). Leptin acts on hypothalamic melanocortin and basal ganglia dopamine receptors to regulate energy homeostasis—and in particular to decrease appetite and inhibit food intake. Fasting-state leptin levels are generally high in patients with obesity before surgery, suggesting resistance to its anorexic action (Stice et al. 2008; Crujeiras et al. 2015), and rapidly decrease after bariatric surgery (to a higher extent than surgery-induced decreases in fat mass) (Faraj et al. 2003). Here we showed sensitivity of the brain’s hedonic valuation system to the drop in fasting-state leptin after RYGB surgery. However, this association does not enable us to causally conclude whether bariatric surgery (through body fat loss) decreased leptin levels, which then act upon dopaminergic projections from midbrain neurons to improve the brain’s hedonic valuation system RSC, or whether improved dopamine functioning is a result of the improved hedonic valuation system’s RSC and independent from the observed decreased leptin levels. Our results open the window for future research investigating the causal links among changes induced by bariatric surgery within the brain’s hedonic valuation system at rest, leptin, and dopamine functioning.

We further found that compared to lean, participants with severe obesity displayed an enhanced RSC between the vmPFC and a set of lateral prefrontal cortex regions that are associated with the cognitive regulation of affective states, working memory and the cognitive control of goal-directed action selection (Ochsner et al. 2002; Wager et al. 2003; Charron et al. 2010). This result is in line with findings from fMRI studies showing an impulse control-related activation of the lateral and dorsolateral prefrontal cortex in patients with obesity (Weygandt et al. 2015). However, investigating connectivity at rest, we did not find a prominent role of vmPFC to dlPFC RSC for weight loss reported in prior task-based fMRI studies (Weygandt et al. 2015).

In summary, our study provides novel evidence that the ability to lose weight is linked to the intrinsic functional organization of the brain’s hedonic valuation system dedicated to reward processing and motivation. We provide evidence that these effects are linked to hormonal homeostatic control that targets hypothalamic and dopaminergic pathways in order to influence food-related behaviour and weight loss. Together, our findings provide a more holistic view between the seldom bridged study of brain systems involved in hedonic aspects of dietary decision-making and its control and homeostatic markers involved in food intake control.

## Acknowledgements

The study was supported by the Sorbonne University IDEX Emergence Grant and ANR ERC-Tremplin Grant (T-ERC CoG) awarded to HP; ICAN research grant funding the leaky gut research awarded to HP, JAW, CPB and KC; and PHRC-(Microbaria) funding awarded to KC. We thank Nicolas Manoharan for collecting the fMRI data; Valentine Lemoine (clinical research assistant, ICAN) for help in clinical investigation; Dr Florence Marchelli (NutriOmics research team) for data management; ICAN CRB members for contribution to bio-banking; Valerie Godefroy for recruiting the control participants; Anne-Dominique Lodeho for technical advice for the rsMRI sequence; Cecile Gallea and Romain Valabrèque for advice in data analysis; Michele Chabert, Armelle Leturque and Patricia Serradas for their feedback during various stages of the project; and Pierre Chandon and Etienne Koechlin for their continuous support in implementing the overall protocol.

## Supplementary information

### Additional clinical assessments of morbidly obese patients before surgery

Obese participants were assessed before bariatric surgery on the following medical exams: depression (BDI, Beck Depression Inventory), alcohol abuse (AUDIT, alcohol use disorders test), nicotine abuse (Fagerstrom), dietary restraint, disinhibition, hunger (TFEQ, three-factor eating questionnaire) and diabetes (clinical assessment). Glycemia was assessed by measuring blood glucose levels after a glucose challenge test (100 ml Fresubine drink) (Table 1) and after overnight fasting (Table S1). Overall, obese participants were not depressed. As shown in supplementary Table 1, on average the obese participant sample was not characterized by alcohol abuse (mean = 6.4, s.e.m. = 1.3, abuse cutoff score ≥ 7, dependence cutoff score ≥ 11); the observed range was between a minimum score of 0 and a maximum score of 12 (*n* = 1 participant). On average, obese participants were not nicotine dependent (mean = 2.3, s.e.m. = 1.3, cutoff score for weak dependence ≥ 4, observed range: 0 to 14 (*n* = 1 obese participant)). Average severity of dietary restraint (mean = 2, s.e.m. = 0.2, observed range: 1 to 3), disinhibition (mean = 1.4, s.e.m. = 0.1, observed range: 1 to 2) and hunger (mean = 1.2, s.e.m. = 0.1, observed range: 1 to 2) was weak to moderate. Blood glucose levels after a glucose challenge test were on average normal in both lean (mean = 4 mmol/l, s.e.m. = 0.1 mmol/l) and obese participants before (6 mmol, s.e.m. = 0.4 mmol/l) and after (mean = 5 mmol/l, s.e.m. = 2 mmol/l) bariatric surgery. However, blood glucose levels in the obese participants sampled after overnight fasting revealed that 80% of the obese participants had glycemia (s.e.m. = 10%, *n* = 6 participants with glucose intolerance, *n* = 7 with type 2 diabetes) compared to 30% after bariatric surgery (s.e.m. = 10%, *n* = 3 with glucose intolerance, *n* = 2 with type 2 diabetes).

### Additional statistical analysis and results

As a robustness check we also calculated residual leptin values by regressing out any variance of leptin explained by kg body fat. We then correlated the difference of before minus after surgery in residual leptin values to the difference in before minus after surgery in vmPFC-vStr RSC, respectively. It revealed a significant covariance (ρ = 0.41, *p* = 0.08, 95% CI due to chance: -0.45–0.46; for % body fat: ρ = 0.52, *p* = 0.05, 95% CI due to chance: -0.45–0.46).

**Table S1:** Clinical assessment of obese patients before bariatric surgery.

|  | Mean | s.e.m. |
| --- | --- | --- |
| Beck Depression Inventory | 1.3 | 0.4 |
| AUDIT | 6.4 | 1.3 |
| Fagerstrom | 2.3 | 1.3 |
| Dietary restraint (TFEQ) | 2.0 | 0.2 |
| Dietary disinhibition (TFEQ) | 1.4 | 0.1 |
| Hunger (TFEQ) | 1.2 | 0.1 |
| % of participants with glycemia before surgery | 0.8 | 0.1 |
| % of participants with glycemia after surgery | 0.3 | 0.1 |

AUDIT: alcohol use disorders test; a score of >7 indicates alcohol abuse, and >11 indicates alcohol dependence in women. Fagerstrom: nicotine dependence questionnaire, 0 to 2 no, 3 to 4 weak, 5 to 6 moderate, 7 to 10 strong nicotine dependence. Severity of dietary restraint, disinhibition and hunger: low = 1, moderate = 2, high = 3. Glycemia reflects the number of participants with glycemia and hyperglycemia.

## Footnotes

* We also tested for a main effect of time but found no differences between T0 and T6 across the whole participant sample, even at a more lenient uncorrected significance threshold of *p* < 0.001.

